# Unsupervised Approach for Electric Signal Separation in Gnathonemus petersii: Linking Behavior and Electrocommunication

**DOI:** 10.1101/2025.03.04.641376

**Authors:** Ivana Chrtkova, Vlastimil Koudelka, Veronika Langova, Jan Hubeny, Petra Horka, Karel Vales, Roman Cmejla, Jiri Horacek

## Abstract

The transfer of information between individuals is fundamental to living systems. Therefore, communication should be studied in various species. Weakly electric fish, *Gnathonemus petersii*, provides a unique model organism for such investigations due to its advanced electrocommunication capabilities, using electric organ discharges (EODs). Separating EODs from multiple individuals is crucial yet challenging. To remediate it, we developed an unsupervised algorithm for EOD separation in two free-swimming individuals. Using continuous wavelet transform, t-distributed Stochastic Neighbor Embedding, and hierarchical clustering, we achieved accurate discrimination of EODs without the necessity of any training data. This approach overcomes the supervised algorithms based on previously published methods in accuracy and computational efficiency, simplifies experimental procedures, and supports animal well-being by reducing the number of required measurements. Additionally, we introduced a novel technique to map electric signals onto auditory representations, facilitating intuitive analysis of EOD sequences. These advancements lay the groundwork for future studies of EOD-based communication, highlighting the potential of *Gnathonemus petersii* in neuroethological, psychopharmacological, and translational research.

## Introduction

Communication is the foundation of information flow, involving exchanging, transmitting, and interpreting signals within a social context. Animal models are essential for studying these processes and understanding the mechanisms underlying information transfer^1–3^. The weakly electric fish *Gnathonemus petersii* (*G. petersii*), capable of emitting and perceiving electric signals, preserves an ideal model for exploring communication principles in complex systems. The fish generates electric organ discharges (EODs) for electrolocation and electrocommunication, enabling navigation and interaction with conspecifics within its environment^4,5^. EODs are produced by an electric organ, located in the fish’s caudal peduncle^6^, which consists of electrocytes whose synchronized excitation generates EODs and establishes a three-dimensional electric field surrounding the fish^7,8^. The waveform characteristics carry information about an individual’s identity^9,10^, while the precise timing of discharges provides insights into the behavioral context and responses to external stimuli^9,11,12^. The analysis of inter-pulse intervals (IPIs) - the time intervals between consecutive EODs - and exploration of generated patterns offer valuable opportunities to elucidate how neural circuits encode and process sensory information, thereby contributing to deciphering the neural code^9^. Furthermore, patterns of EOD rhythms may provide novel approaches to modeling language-related symptoms of mental disorders, positioning *G. petersii* as a promising animal model for translational research^13,14^ to remediate the pitfalls of the standard animal models^15,16^.

EOD of *G. petersii* is characterized by a biphasic pulse pattern^17,18^ with a brief duration of approximately 200-400 μs^19,20^. The short pulse length presents a considerable challenge in developing classifiers capable of accurately separating EODs from multiple individuals recorded simultaneously. The frequency of EOD generation depends on the behavioral context and may exceed 100 Hz^20,21^. In social experiments, recordings often capture thousands of EODs from multiple individuals, making manual separation impractical and underscoring the need for automated algorithms to enhance the efficiency of signal processing.

With the advancement of machine learning methods, various sophisticated algorithms that solve this task with high accuracy have been developed^22–24^. However, these methods typically rely on supervised learning and require pre-recorded EODs from a single fish to create a training dataset for assigning EODs to corresponding individuals from recordings of dyads. A simple automated method (that does not require manual assignment) is based on the cross-correlation of EODs from dyad recordings with templates obtained during the recordings of a single fish in the aquarium^25^. However, this approach is insufficient when individuals exhibit similar EOD durations or discharge almost at the same time. More advanced algorithms include the classification with support vector machine (SVM) utilizing extracted features from EODs^22,23^ or combining the waveform characteristics and positional information from video tracking data^24^. Although effective, these methods often require substantial computational resources or complex experimental setups. Additionally, the fish must undergo several measurements to obtain training data for classifiers, which may be stressful for individuals involved in the experiment. The accuracy of classification may also be limited since the training data originates from different recording sessions, introducing additional variability that cannot be explained by the model.

To remedy these pitfalls, our study aimed to develop a novel algorithm based on unsupervised learning for separating EODs from two free-swimming simultaneously recorded *G. petersii* fish. Before constructing the classifier, the inter-individual variability in EOD waveforms was explored to determine the most effective feature representations for capturing this diversity and achieving the highest classification accuracy. The performance of the developed algorithm was compared with the classification results from the two algorithms based on supervised learning to investigate the effectiveness of the unsupervised approach. Finally, we introduced a novel technique to link electric activity with behavior through sonification, enabling perceptual exploration of generated EOD patterns.

## Results

### Exploring inter-individual variability of EODs

The first step in our analysis was to explore inter-individual variability in the EOD waveforms across 24 individuals recorded separately (Dataset 1). As shown in **Figure 1a-c**, the 2D mappings produced by the t-SNE algorithm using three different EOD representations - in the time domain, frequency domain, and time-frequency domain - demonstrate significant differences in cluster compactness. t-SNE was run with perplexity values ranging from 30 to 120, and the final output was selected based on visual inspection to ensure the most distinguishable separation of clusters. The time-frequency domain representation, obtained by continuous wavelet transform (CWT), resulted in the most distinct and well-separated clusters, highlighting its effectiveness in capturing inter-individual variability compared to the time and frequency domain representations. However, a downside of using time-frequency domain features was the significantly longer computation time for the t-SNE algorithm (907.82 seconds), compared to the frequency domain (241.22 seconds) and the time domain (176.89 seconds) representations.

**Figure 1:**
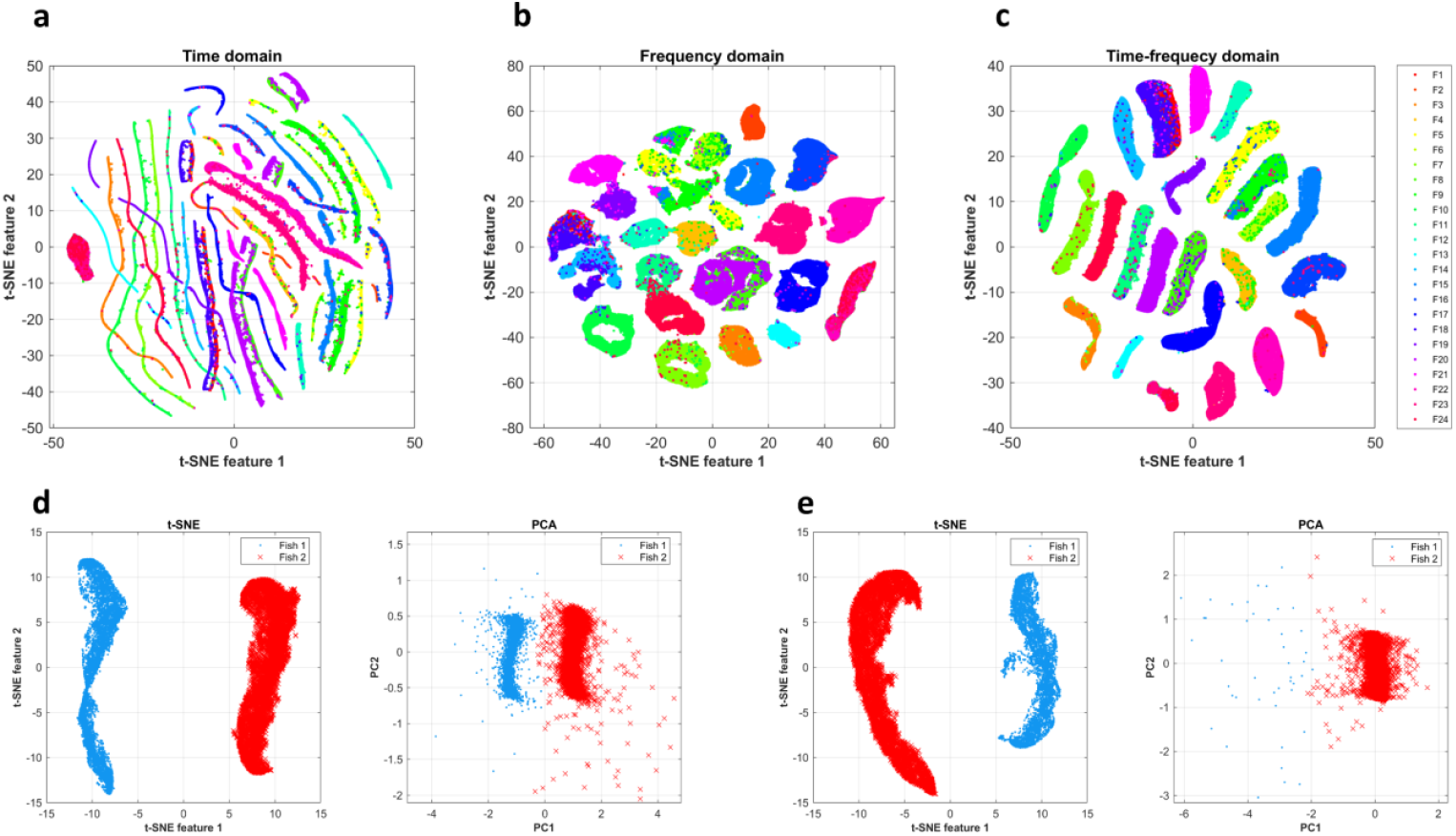
Exploration of inter-individual variability with dimensionality reduction techniques. **a-c**, Images illustrate outputs from the t-SNE algorithm with EOD features extracted from different domains using Dataset 1. That is: **a**, original waveforms in the time domain, **b**, Fourier transform of original waveforms in the frequency domain, and **c**, coefficients of continuous wavelet transform in the time-frequency domain. A different color label is used for each of the 24 individuals from Dataset 1. **d-e**, Comparison of the visualizations of time-frequency features utilizing two dimensionality reduction techniques: t-SNE (left) and PCA (right) on recordings from Dataset 2. **d**, Illustration of a scenario where two distinct clusters can still be observed after applying PCA. **e**, Illustration of a scenario where the t-SNE algorithm effectively identifies two easily separable clusters, while PCA suggests the data are not easily separable into two classes. The distribution of these two cases was balanced at 50/50.

### Comparison of linear and nonlinear dimensionality reduction methods

In the analysis of the dyad’s recordings from Dataset 2, time-frequency features of EODs were extracted and subsequently visualized in a two-dimensional space using t-SNE and PCA. PCA revealed two distinct clusters in 50% of the recordings; however, in the other half, only a single cluster was observed and hierarchical clustering did not successfully categorize the fish into two groups. In contrast, t-SNE consistently identified distinct clusters in each instance. **Figure 1d-e** illustrates two types of results generated by t-SNE and PCA alongside the classifications produced by hierarchical clustering.

### Performance of algorithms

Table 1. summarizes the median values and interquartile ranges of performance metrics for all algorithms applied to 10 recordings of dyads. Our developed algorithm based on t-SNE and hierarchical clustering was executed with consistent hyperparameter setting, always identifying two distinct and well-separated clusters. It outperformed the correlation method (ACC = 0.688, MCC = 0.472) and the SVM classifier (ACC = 0.898, MCC = 0.747), achieving an accuracy of 0.993 and a Matthews correlation coefficient of 0.981, with classification completed in approximately 35 seconds per recording.

**Table 1:** Results from validation of classifiers’ performance. Median values and interquartile ranges (in round brackets) of accuracy (ACC), Matthews correlation coefficient (MCC), and running time (RT) for all employed separation algorithms. Values were calculated based on the performance outcomes from 10 recordings of dyads (Dataset 2).

| Algorithm | ACC (-) | MCC (-) | RT (s) |
| --- | --- | --- | --- |
| t-SNE + HC | 0.993 (0.007) | 0.981 (0.015) | 34.978 (15.057) |
| Correlation method | 0.688 (0.311) | 0.472 (0.333) | 2.794 (1.671) |
| SVM | 0.898 (0.186) | 0.747 (0.387) | 1267.341 (536.747) |

**Figure 2a** shows the distribution of performance metrics in the form of boxplots. The supervised classifiers achieved a high value of accuracy and Matthews correlation coefficient on some recordings, while on others, their performance was more similar to that of a random classifier. In comparison, our algorithm achieved high values of performance metrics with low interquartile ranges, indicating consistent performance across the recordings. The effectiveness of our algorithm based on t-SNE and hierarchical clustering is demonstrated in **Figure 2c**. On a randomly selected segment, our algorithm correctly classified all EODs, while the other two algorithms showed some misclassifications.

**Figure 2:**
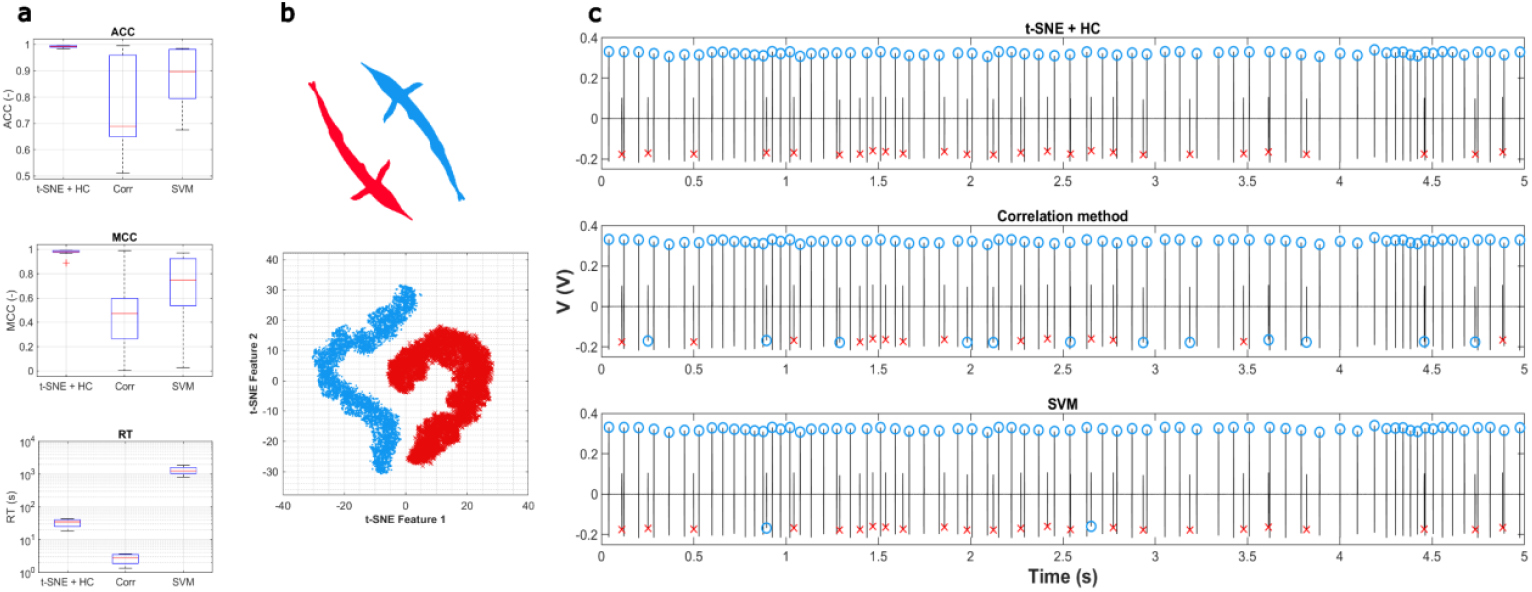
Comparison of classifiers’ performance. **a**, Boxplots depicting values of performance metrics (from top: Accuracy, Matthews correlation coefficient, and Running time) obtained from Dataset 2. The reason for displaying the running time (RT) metric with a logarithmic scale on the y-axis is the desired visibility of differences between algorithms, which would not be evident in a linear scale due to the long duration of the SVM classifiers’ training phases. Boxplots illustrate the highest ACC and MCC median values with small interquartile ranges for the t-SNE + HC algorithm. **b**, Result of 2D mapping of feature vectors from a randomly selected recording of dyads obtained using the t-SNE algorithm. Clusters are colored with classification labels from hierarchical clustering corresponding to two individuals. **c**, Comparison of the results from a classification using (from top) t-SNE + HC algorithm, correlation method, and SVM classifier on the 5-second segment from the same electric signal as in **b**. Color markers on EODs correspond to clusters color in **a**. Only the t-SNE + HC algorithm correctly assigned all EODs on the illustrated signal segment.

Due to the large interquartile ranges observed in the performance of supervised classifiers, we aimed to explore the dependency of supervised classifiers’ performance on differences in the measured body characteristics of the fish in dyads. No relationship was confirmed, as the correlation analysis did not reveal any significant findings. The summary of results from the correlation analysis is presented in **Table 2**.

**Table 2:** Results from correlation analysis to explore dependency of classification outcomes on differences in body characteristics. Values of Pearson correlation coefficient and corresponding *p*-values between the results of performance metrics (accuracy, Matthews correlation coefficient) of two supervised algorithms and differences in body parameters of fish in a measured dyad. No significant relationship between the algorithms’ performance and differences in physical characteristics was observed.

| | | $\Delta$ Total weight | | $\Delta$ Total length | | $\Delta$ Standard length | |
| --- | --- | --- | --- | --- | --- | --- | --- |
|  |  | ACC | MCC | ACC | MCC | ACC | MCC |
| Correlation method | R | 0.0868 | 0.0995 | -0.4767 | -0.415 | -0.267 | -0.2095 |
|  | p | 0.812 | 0.7844 | 0.1636 | 0.234 | 0.456 | 0.561 |
| SVM | R | 0.177 | 0.186 | 0.0674 | -0.0364 | -0.0461 | 0.1624 |
|  | p | 0.626 | 0.607 | 0.853 | 0.921 | 0.899 | 0.654 |

### Sonification of EOD signal

EODs from the long recording of the dyad (Dataset 3) were classified with our developed unsupervised algorithm, and the signals were sonified and integrated into the video recording. The demonstrations of these audio-visual representations are provided as supplementary materials. The first example consists of two videos (**S1** and **S2**), each containing the sonified signal from one of the two recording channels with preserved information about the amplitude of EODs (pulse-wise approach). This example illustrates how this approach can serve as an additional method for validating the separation process since it demonstrates how the electric signals of both fish amplify or attenuate as they approach the recording channels or move away from them. The remaining video examples (**S3** and **S4**) highlight the potential of the sonification techniques in studying fish interactions, providing a novel method to link electrocommunication with behavioral dynamics. In video **S3**, the sonification allows us to hear fluctuations in EOD rates while maintaining the precise temporal alignment of individual EODs, making it possible to perceive how the timing of electrical discharges changes during interactions. On the other hand, video **S4** facilitates the examination of variations in EOD rates through evolving melodies, which may give rise to both, harmonic and dissonant structures, providing deeper insight into the complex dynamics underlying electrocommunication. We also provide complete audio tracks (**S5** and **S6**), enabling a more comprehensive exploration of EOD interactions themselves over longer temporal scales. Supplementary materials are available on Zenodo (https://doi.org/10.5281/zenodo.14960395).

## Discussion

The main result of this study is an effective method for capturing inter-individual variability in EOD shapes by applying continuous wavelet transform for feature extraction and t-SNE for dimensionality reduction. It is essential for the possibility of a detailed study of the electrocommunication (or even proto-language) of this unique species. The time-frequency representation using scalograms enabled a more accurate differentiation between the 24 individuals recorded separately, compared to the original waveforms in the time domain or their frequency spectra obtained via fast Fourier transform. This was evident from the highly compact and easily distinguishable clusters observed (**Figure 1a-c**). Our findings extend previous research focused on the classification between sympatric species of the weakly electric fish *Gymnotus*^26^ by showing that the superiority of time-frequency representation is also observable within the intraspecies identification. The combination of scalograms and t-SNE proved to be a powerful approach in achieving the high classification performance demonstrated in our study. Although this method yields high accuracy, it requires extended processing time due to the many wavelet coefficients used as inputs to t-SNE.

The performance metrics results demonstrate (**Table 1, Figure 2a**) that our unsupervised algorithm, which incorporates hierarchical clustering for classifying the 2D mapping obtained from t-SNE, outperformed the correlation method and SVM classifier. One of the key advantages of this method is that it eliminates the necessity of recording individual fish separately to create a training dataset, thereby reducing the overall number of recordings needed for the experiment and offering an alternative approach to existing supervised algorithms for EOD separation^22–25^. This simplification makes the experimental procedure more efficient and primarily, minimizes stress for the animals involved. Also, we avoided the time-consuming training phase of the SVM or any other advanced supervised algorithm that could be potentially used for classification. Additionally, the algorithm generates visual representations of high-dimensional data, which can serve as an initial indication of successful separation. For instance, the presence of two distinct clusters in the 2D mapping may suggest that the algorithm correctly distinguished the signals from two fish, providing a useful first step in the validation process.

It is important to note that both supervised algorithms exhibited large interquartile ranges in the performance metrics examined (**Table 1, Figure 2a**). While the algorithms demonstrated high performance on some recordings, their behavior was similar to random classifiers on others, suggesting that the classification accuracy of supervised algorithms depends on the specific combination of individuals. The observation is further supported by the finding that PCA revealed two clusters in half of the cases and one cluster in the other half (**Figure 1d,e**)). For this reason, we explored potential correlations between the differences in measured body characteristics of recorded fish and classification outcomes, but no significant relationship was found (**Table 2**). Furthermore, there were no relations between the two types of PCA behavior and the classifiers’ performance or differences in body parameters. Therefore, the reason for the high variability in the separation success/failure remains unclear, and the following research may focus on the exploration of these differences.

In future studies, the dependency of variations in EOD waveforms on the variability of fish phenotypes should be examined in more detail. The analysis should certainly account for the influence of the individuals’ sex since the shape of EOD is influenced by this factor^27,28^. EOD characteristics may also be modulated by dominant-like behavior, as the shape of the EOD may serve as an indicator of social status^29^. Our algorithm provides a powerful tool for systematically exploring these effects and their implications for classification accuracy.

Given its exceptional electrocommunication abilities, *G. petersii* may represent a unique model organism for studying neural mechanisms underlying communication deficits, particularly in psychiatric disorders associated with speech impairments^14,16,30^. Future studies could reveal how pharmacological manipulations alter the microscopic structure of EOD waveforms, as psychoactive substances such as ketamine influence behavioral patterns and the macroscopic structure of electric signals (e.g. number of generated EODs in single fish)^14^. Most importantly, exploration of EOD communication patterns could provide insights into social interaction deficits observed in neuropsychiatric conditions.

One of the limitations of our proposed algorithm is the need for manual assignment of separated signals to the corresponding individual. Although the algorithm effectively distinguishes two clusters, it does not assign each cluster to a specific fish. This process is relatively straightforward, as the polarity of the EODs is influenced by the fish’s position in the aquarium, allowing accurate identification. However, the processing pipeline still requires user intervention. In further development, incorporating fish tracking data into the classification process^24,31^ could improve accuracy and eliminate the need for manual assignment, thus enabling a fully automated procedure for signal separation. Another limitation of our approach is its suitability only for offline analysis since it is not designed for real-time processing.

Our proposed technique of integrating electric activity into the video recordings through sonification could be used as an additional approach for validating the separation process but primarily as a tool for linking patterns in electric activity with behavior dynamics to explore their underlying connection. The interactions through EOD sequences can be observed and correlated with fish behavior, which sets an initial step for researchers in establishing an EOD-behavior state space to explore communication and behavior, both in basic research of weakly electric and in preclinical modeling of psychiatric diseases.

The ability to accurately separate EOD signals without prior training data provides a significant advancement in electrocommunication research, serving as an initial step toward studying social interactions in weakly electric fish. The integration of sonification techniques offers a novel way to interpret complex electric signal patterns, enabling a more intuitive analysis of EOD sequences and their temporal dynamics. With these methodological improvements, future electrocommunication studies can be significantly enhanced, facilitating deeper insights into the analysis of social behavior and establishing *G. petersii* as a novel animal model in neuroscience.

## Methods

### Animals

A total of 31 fish of the *G. petersii* species were obtained from a local distributor, VIVARIUM Melnik (Melnik, Czech Republic). The sex of the individuals was not determined since sexual dimorphism in *G. petersii* manifesting by alteration in EODs is season-dependent and prolonged captivity can reverse these differences^27,28^. The 126 L experimental aquarium was refilled with 30 L of fresh water before each experiment. The water conductivity was 272.32 ± 22.01 μS, temperature 24.74 ± 0.54 °C, and pH 7.4 ± 0.21. The experimental aquarium was illuminated with red light with an intensity of 9 lux on the water surface, due to the high sensitivity of the cone cells of *G. petersii* to deep-red wavelengths, simulating the red-dominated turbid waters characteristic of its natural habitat^32,33^.

### Experimental design

Three different datasets were used in this study. Dataset 1 was acquired by recording 24 individuals (total weight = 10.37 ± 2.76 g, standard length = 7.85 ± 1.24 cm, total length = 12.30 ± 1.13 cm) separately to obtain 15-minute recordings. This dataset was used to explore the variability in EOD shapes between individuals and to discriminate the most effective representation of EODs based on these findings. Dataset 2 was used to develop and validate the most appropriate classifier for separating signals from two free-swimming individuals. Five individuals (total weight = 7.08 ± 1.46 g, standard length = 7.06 ± 0.62 cm, total length = 11.37 ± 0.73 cm) were first recorded for 15 minutes separately to obtain training data for the supervised classifiers (first part). Subsequently, all possible combinations of pairs were recorded, resulting in 10 recordings of dyads (second part). Six minutes of each dyad recording were extracted to evaluate the classifiers’ performance. Dataset 3 comprises a single long dyad recording (45 minutes), acquired for the EOD classification and demonstration of a new approach for exploring patterns in electric activity through sonification.

### EOD recording

EODs were recorded using the same experimental setup established in our previous studies^14,30^. A specialized data acquisition system comprising three hardware layers: sensor electrodes, an amplifier, and a data acquisition unit. The first layer consisted of Ag electrodes, originally designed for electroencephalographic recording, with four active bipolar-connected electrodes positioned at each corner of the experimental aquarium and a fifth reference electrode in its center. All electrodes were submerged 2 cm below the water surface. Subsequently, the signal was amplified using an instrumental amplifier with a gain of 10. The amplified output was directed to a National Instruments USB-6003 DAQ unit with a sampling rate of 50,000 samples per second and 16-bit resolution for each channel. Data were displayed and stored on a computer with a National Instruments application. Illustration of the experimental aperture, data acquisition and subsequent processing of recorded EODs is depicted in **Figure 3**. The entire acquisition system was synchronized with the 1.3 MPx infrared recording camera (IDS Imaging Development Systems GmbH, Germany) using a common clock cycle to enable accurate synchronization of EOD signal acquisition with digital image capture from an infrared camera.

**Figure 3:**
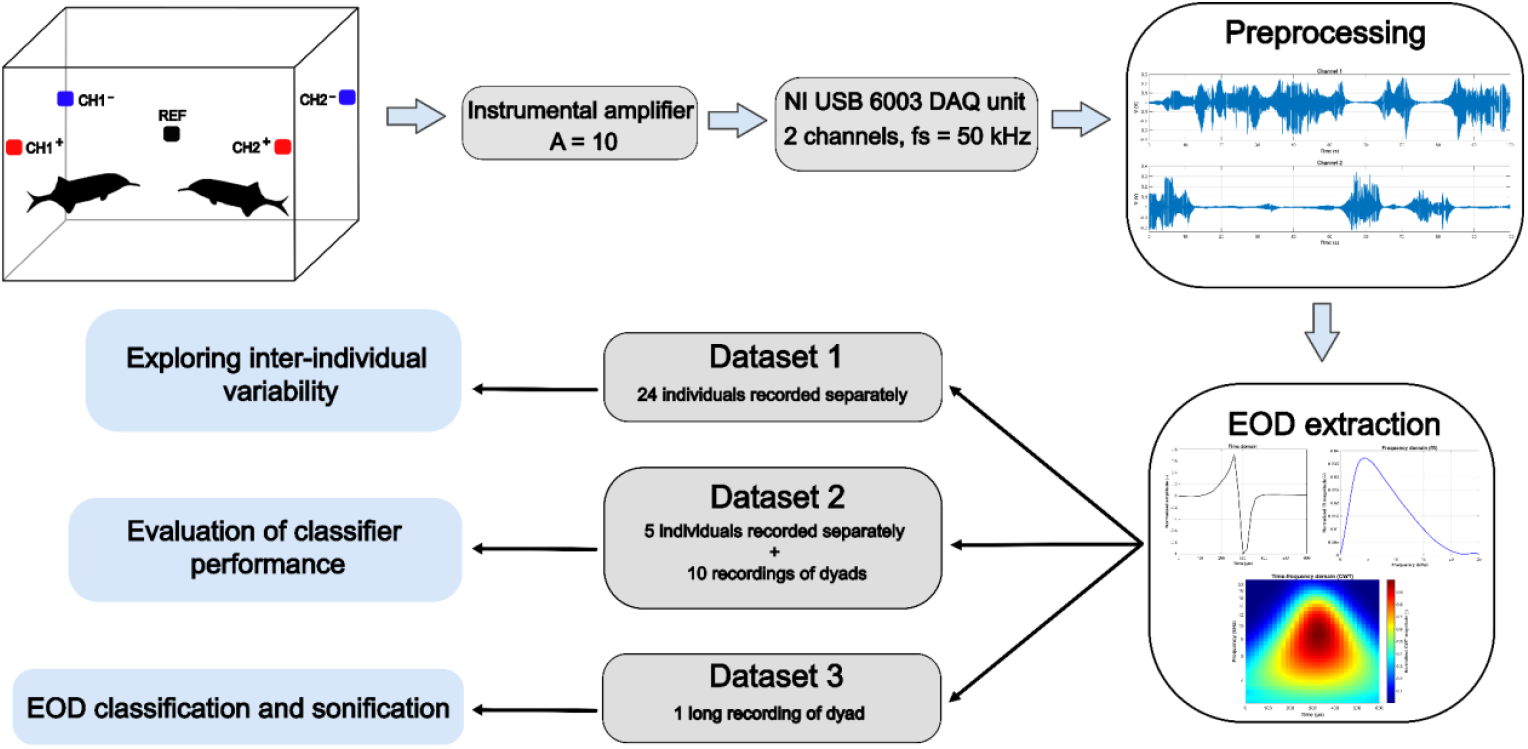
Diagram of recording setup and processing pipeline used in this study. Two channels were recorded with four active bipolar-connected electrodes and one electrode as a reference. Recorded signals were amplified and sampled with the sampling frequency of 50 kHz for each channel. Digitized signals were preprocessed and individual EODs were extracted utilizing 300 μs before and after the detected EOD and stored in the form of normalized waveforms in the time domain, normalized spectra of original waveforms using fast Fourier transform, and normalized scalograms obtained from continuous wavelet transform. Signals from three different datasets were used for exploration of intra-individual variability (Dataset 1), the construction and validation of separation algorithms: Correlation method, SVM, and t-SNE + hierarchical clustering (Dataset 2), and the classification and sonification of EODs (Dataset 3).

### EOD preprocessing and extraction

Recorded signals were analyzed using MATLAB 2023b. The signal isolines approximated through median filtering were subtracted from the signals to eliminate the baseline drift. Before applying the EOD detection algorithm, the signal was squared to enhance the peak amplitude and ensure, that the algorithm accurately identifies the maximum peak of the waveform since the EOD polarity is influenced by the orientation of the fish relative to the electrodes. EODs were detected using the threshold of 5 mV and a minimum distance of 400 μs between peaks. The threshold for EOD detection was determined based on the estimated noise level from experimental recordings (RMS_noise_ = 0.2 mV). EOD detection was performed on both channels; however, only the EOD with higher amplitude was considered for the subsequent analysis. EODs were extracted from the signal utilizing 300 μs before and after the absolute maximum peak of the waveform. Since the amplitude of the EOD depends on the size of individuals^34^ and their position in the aquarium, the EODs were normalized by the value of absolute maximum peak (voltage range of ± 1V) and eventually stored with reversed polarity to maintain consistency between the orientation of the waveforms.

### EOD representations

It was necessary to find the appropriate representation of the EODs capturing the highest inter-individual variability. According to Crampton et al.^26^, using time-frequency features facilitates the classification of different species of weakly electric fish. For this reason, we applied continuous wavelet transform (CWT) with Morse wavelet (γ = 3, P^2^ = 60, 12 Voices per Octave) to each extracted EOD. The mentioned parameters were selected as they offer a sufficient trade-off between time and frequency resolution. Before the transformation, the EODs were padded with zeros (15 zeros on each side), to achieve higher frequency resolution and eliminate boundary effects. After the transformation, scalograms were normalized by the coefficient with maximal value and cropped back to the size of 31 samples (∼620 μs). The resulting coefficients were concatenated into a single feature vector. To confirm the efficiency of time-frequency features, the inter-individual differences in EOD shapes were also examined in the time domain - using original normalized EODs and in the frequency domain - using normalized spectra of EODs obtained with fast Fourier transform. All the representations utilized for the analysis of inter-individual variability are illustrated in **Figure 4**. A total of 235,664 EODs from all 24 individuals in Dataset 1 were extracted and stored in three separate matrices utilizing the described representations (time, frequency, and time-frequency domains).

**Figure 4:**
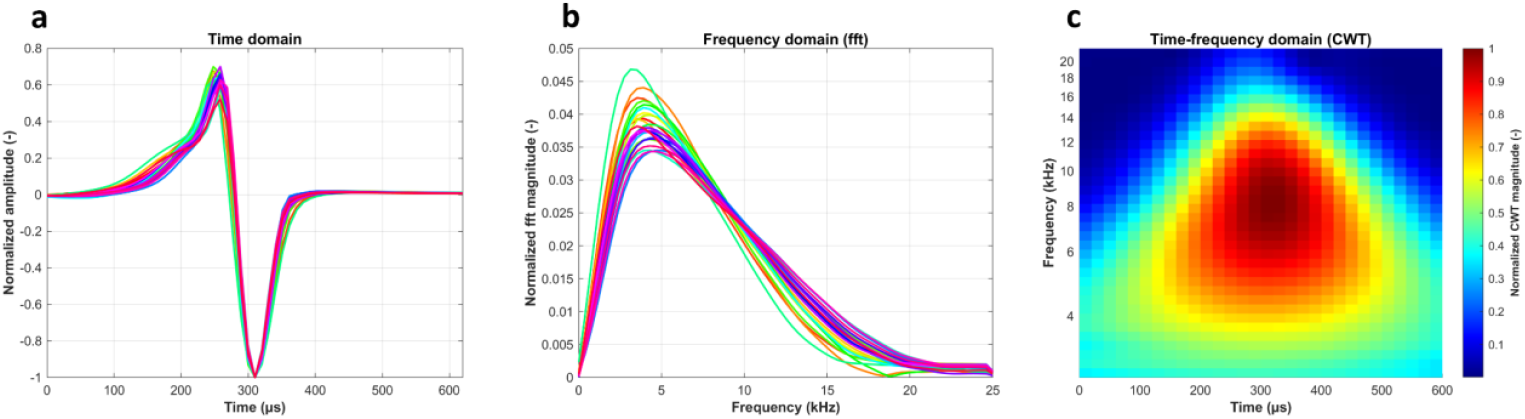
Illustration of three different representations of EODs for exploration of inter-individual variability. **a**, Averaged EODs for 24 individuals from Dataset 1 in the time domain (interpolated for visualization), **b**, frequency domain - obtained by fast Fourier transform of averaged EODs, and **c**, time-frequency domain - obtained by continuous wavelet transform of grand average from 24 individuals. In **a** and **b**, each individual is represented by a different color label, with colors assigned in accordance with **Figure 1a-c**.

### Dimensionality reduction

Since our methodology operates with high-dimensional features of EODs, it is essential to reduce the dimensionality of the data with a suitable dimensionality reduction method before classification. Employing linear techniques for projection into low-dimensional space may be insufficient when dealing with data lying on nonlinear manifolds. In such instances, the application of nonlinear dimensionality reduction methods is more suitable, as these approaches are better equipped to capture the global structure of the data. Among these methods, the t-Distributed Stochastic Neighbor Embedding (t-SNE) algorithm is primarily aimed to visualize high-dimensional data^35^. To this end, the t-SNE was employed to explore inter-individual variability using Dataset 1 and as an initial step for an unsupervised classification algorithm for EODs separation from Dataset 2. The open-source implementation of t-SNE, incorporating polynomial interpolation and fast Fourier transform^36^ was applied to enhance time efficiency.

Results from the t-SNE algorithm obtained from Dataset 2 were compared by visual inspection with the outputs using principal component analysis (PCA), a widely utilized linear dimensionality reduction method. This technique provides an efficient and well-interpretable solution; however, it may not sufficiently uncover the underlying structure of the data.

### Classification

The 2D mappings of recordings from Dataset 2, obtained by the t-SNE algorithm, were classified into two clusters using hierarchical clustering with the average linkage method and Euclidean distance. Hierarchical clustering was selected due to the presence of non-linearly separable clusters after dimensionality reduction, making linear decision boundary-based algorithms, such as k-means, unsuitable for this task. The novel unsupervised approach did not require any prerequisites and training data based on the first part of Dataset 2.

The correlation method and support vector machines (SVM) classifier with radial basis function (RBF) kernel were employed as two supervised approaches for comparison with our developed unsupervised algorithm. For the correlation method, the average scalograms from the recordings of a single fish in Dataset 2 (the first part) were obtained for each of the two corresponding individuals in the dyad recording in Dataset 2 (the second part). These averaged scalograms were used as templates for the computation of correlation coefficients with scalograms extracted from dyad recording. Then, the EOD from dyad recordings was assigned to the individual with the higher Pearson’s correlation coefficient value.

For the training phase of the SVM classifier, 5,000 EODs with the highest signal-to-noise ratio, determined by the amplitude of the waveform in the time domain, were extracted from the single fish recordings in the first part of Dataset 2. Subsequently, wavelet coefficient predictors were standardized by the corresponding weighted column mean and standard deviation. For each recorded dyad, a model was trained to recognize the two corresponding fish in the second part of Dataset 2. Hyperparameters of the SVM RBF kernel (γ, C) were optimized using Bayesian optimization with 10-fold cross-validation.

### Performance evaluation

To evaluate the performance of the classifiers, EODs from all recordings of dyads were manually assigned to the corresponding individual based on the amplitude and polarity of the signal relative to the individual’s position within the aquarium. A total of 118,050 EODs were assigned in this manner to create the test dataset. The procedure was performed twice to ensure the reliability of the manual assignment. Although there was a disagreement in approximately 0.5% of the instances, the differences in performance metrics of the algorithms clearly exceeded this level of uncertainty. Therefore, we re-evaluated the ambiguous cases and proceeded with our analysis.

The algorithms were evaluated based on the accuracy (ACC), the Matthews correlation coefficient (MCC), and running time (RT). Given the significant variability in the counts of EODs during social interactions, the Matthews correlation coefficient was selected as a performance metric due to its robustness when applied to unbalanced datasets^37^. The MCC ranges from -1 to 1, with a value of 1 indicating a perfect classification, a value of -1 representing a complete misclassification of all samples, and a value of 0 signifying a random prediction. The duration of the training phase of the SVM algorithm was included in the running time value.

After quantification of the algorithms’ performance, Pearson correlation coefficients between the performance metrics and differences in measured body characteristics (total weight, total length, standard length) of recorded individuals in dyad were computed to explore the potential relationship between the classification outcomes and variations in fish’s physical parameters.

### EOD sonification for application in research

For a study of electrocommunication and exploration of relationships between patterns in electric activity and behavior, it is beneficial to have the capability to integrate electric signals into video recordings. An efficient way to address this task is to create sonified signals and incorporate them into the video as a sound component. Due to the short duration of EODs and relatively small inter-individual variability, it would not be possible for the human ear to distinguish two individuals solely based on their sonified EODs. To remediate this point, sinusoidal waveforms with different frequencies for each individual (instead of original EODs) were inserted at the corresponding instances of original EODs generated by fish. The duration of the sinusoidal waveforms was preserved the same as the duration of extracted EODs in previous analyses (620 μs) and amplitudes of the waveforms were weighted by amplitudes of original EODs if maintaining information about the original signal intensity was desired.

Sonified sinusoidal waveforms with such a brief duration are perceived as auditory clicks rather than pure tones, as the pitch of sinusoidal tones is influenced not only by their fundamental frequency, but also by other factors including duration, intensity, and amplitude envelope^38–41^. Due to this perceptual effect, it was necessary to determine an appropriate combination of frequencies to ensure clear distinguishability between the two sonified waveforms despite their short duration. Experimental testing revealed that a combination of 1.5 and 15 kHz resulted in the most distinct auditory discrimination between the two individuals while preserving the temporal alignment of EODs. Following the separation of EODs from Dataset 3, the signals were sonified as described and integrated into the video recordings. Created videos also serve as an additional form of validation of the performance of separation algorithms.

To explore the structure of EOD signals in two interacting fish, frequency modulation (FM) sound synthesis was applied. First, two distinct and harmonious carrier frequencies, *fc1* and *fc2*, were selected to serve as specific frequency codes for each fish, similar to the approach described earlier: *fc1* was set to 523.3 Hz and 1046.5 Hz (tones C5 and C6), and *fc2* to 392 Hz and 784 Hz (tones G4 and G5). Instead of a pulse-wise approach, continuous sonification of the two harmonic signals was used.

To enhance auditory readability, both harmonic signals were extended by their second harmonics (C5 by C6 and G4 by G5). The carrier frequencies were continuously modulated based on the time interval between two consecutive EODs (IPI) of the fish. Notably, the IPI time series was low-pass filtered using a Butterworth filter with a cutoff frequency of 1 Hz before modulation, ensuring smooth variations in the distinct harmonious frequencies corresponding to different EOD rates. The filtered IPI time series sign was then inverted to provide a direct proportion between the EOD frequency and the carrier frequency. This process mapped the temporal evolution of the two EOD rates, representing the two interacting fish, onto two distinct melodies.

The degree of coupling between the signals, in terms of EOD rates, could be observed as the level of independence between the two melodies. Furthermore, since the two carrier frequencies were chosen to be harmonious, the harmony or disharmony between the two “voices” also played a role. Harmonic states represent situations where the EOD signals of the fish are in perfect synchrony, whereas disharmonic states indicate a lack of synchrony.

## Data and code availability statement

MATLAB codes for the developed separation classifier and both EOD sonification approaches are freely available on GitHub (https://github.com/ivanachrtkova/EOD_tools). The raw data supporting this article’s conclusions will be made available upon request without undue reservation.

## Ethics statement

The animal study was approved by the ethics committee of Charles University (registration number 19014/2019-MZE-18134, MSMT approval number 27367/2019-3) and was in accordance with the local legislation and institutional requirements.

## Acknowledgements

The present study was supported by the Czech Health Research Council (grant number AZV CR NU21 04–00405, NW24-04-00413 and NW25-04-00454), Horizon Europe (grant no. 101137378, HORIZON-HLTH-2023-DISEASE-03-01) and ERDF-Project Brain dynamics, No. CZ.02.01.01/00/22 008/0004643 and VVI CZECRIN (LM2023049).

## Author contributions

**IC**: Paper structure, Software, Validation, Writing - original draft preparation; **VK**: Supervision, Writing - original draft preparation; **VL**: Methodology, Data acquisition, Writing - review and editing; **JaH**: Data curation, Preprocessing; **PH**: Funding acquisition, Providing expertise in fish experiments; **KV**: Project administration, Funding acquisition, Providing expertise in translational research; **RC**: Providing expertise in signal processing, Writing - review and editing; **JiH**: Funding acquisition, Methodology, Providing expertise in animal models of psychosis, Writing - review and editing;

